# A non-canonical RNAi pathway controls virulence and genome stability in Mucorales

**DOI:** 10.1101/2020.01.14.906289

**Authors:** C. Pérez-Arques, M.I. Navarro-Mendoza, L. Murcia, E. Navarro, V. Garre, F. Nicolás

**Author notes:** Equally contributed (C. Pérez-Arques, M.I. Navarro-Mendoza).

## Abstract

Epimutations in fungal pathogens are emerging as novel phenomena that could explain the fast-developing resistance to antifungal drugs and other stresses. These epimutations are generated by RNA interference (RNAi) mechanisms that transiently silence specific genes to overcome stressful stimuli. The early-diverging fungus *Mucor circinelloides* exercises a fine control over two interacting RNAi pathways to produce epimutants: the canonical RNAi pathway and a new RNAi degradative pathway. The latter is considered a non-canonical RNAi pathway (NCRIP) because it relies on RNA-dependent RNA polymerases (RdRPs) and a novel ribonuclease III-like named R3B2 to degrade target transcripts. Here in this work, we uncovered the role of NCRIP in regulating virulence processes and transposon movements through key components of the pathway, RdRP1, and R3B2. Mutants in these genes are unable to launch a proper virulence response to macrophage phagocytosis, resulting in a decreased virulence potential. The transcriptomic profile of *rdrp1*Δ and *r3b2*Δ mutants revealed a pre-exposure adaptation to the stressful phagosomal environment even when the strains are not confronted by macrophages. These results suggest that NCRIP represses key targets during regular growth and release its control when the fungus is challenged by a stressful environment. NCRIP interacts with the RNAi canonical core to protect genome stability by controlling the expression of centromeric retrotransposable elements. In the absence of NCRIP, these retrotransposons are robustly repressed by the canonical RNAi machinery; thus, supporting the antagonistic role of NCRIP in containing the epimutational pathway. Both interacting RNAi pathways might be essential to govern host-pathogen interactions through transient adaptations, contributing to the unique traits of the emerging infection mucormycosis.

**Author summary:** Mucormycosis is an emergent and lethal infectious disease caused by Mucorales, a fungal group resistant to most antifungal drugs. *Mucor circinelloides*, a genetic model to characterize this infection, can develop drug resistance via RNAi epimutations. This epimutational RNAi mechanism interacts with a novel non-canonical RNAi pathway (NCRIP), where the ribonuclease III-like R3B2 and the RNA-dependent RNA polymerase RdRP1 are essential. The analysis of the transcriptomic response to phagocytosis by macrophage in *rdrp1*Δ and *r3b2*Δ mutants revealed that NCRIP might control virulence in *M. circinelloides*. These mutants showed constitutive activation of the response to phagocytosis and a reduction in virulence in a mouse model, probably caused by a disorganized execution of the genetic program to overcome host defense mechanisms. The antagonistic role of the NCRIP and the RNAi canonical core is evident during post-transcriptional regulation of centromeric retrotransposons. These retrotransposons are silenced by the canonical RNAi pathway, but this regulation is restrained by NCRIP, proven by an overproduction of small RNAs targeting these loci in NCRIP mutants. These new insights into the initial phase of mucormycosis and transposable element regulation point to NCRIP as a crucial genetic regulator of pathogenesis-related molecular processes that could serve as pharmacological targets.

## Introduction

Mucorales are a group of ancient fungi that are emerging as a new source of pathogens causing the fungal infection mucormycosis. This infectious disease is increasing the focus of recent studies due to its high mortality rates, which can reach up to 90% in cases of disseminated infection [1,2]. The elevated mortality rate is a direct connection to a lack of effective antifungal treatments, a consequence of the unusual resistance observed in these fungi. In this regard, a novel RNAi-dependent epimutational mechanism of drug resistance has been described in *M. circinelloides* [3]. In this mechanism, *M. circinelloides* generates strains resistant to the antifungal drug FK506 after only four days of exposure. The mechanism behind this rapid adaptation relies on the specific silencing of the *fkbA* gene and its encoded FKBP12 protein, which is the target of FK506. Thus, in the absence of FKBP12 due to *fkbA* silencing, the drug FK506 is unable to hinder the mycelial growth of *M. circinelloides*, generating transient resistant strains that arise due to selective pressure. The epimutational drug resistance in *M. circinelloides* is becoming clinically relevant because epimutants can emerge upon exposure to other antifungal drugs [4], and they exhibit organ-specific stability during in vivo infection [5].

The RNAi pathway involved in this epimutation-based drug resistance depends on the canonical components of the RNAi machinery, which are broadly characterized in *M. circinelloides* [6]. First, RNA dependent RNA polymerases (RdRPs) generate double-stranded RNA (dsRNA). Later, dsRNA is processed by RNase III Dicer enzymes to generate small RNAs (sRNAs). Then, the third element of the RNAi canonical core, the Argonaute protein (Ago), uses the sRNAs to conduct homology-dependent repression of the target sequences [7]. Besides drug-resistance, the canonical core elements participate in RNAi-based defensive pathways protecting genomic integrity against invasive nucleic acids and transposable elements, as well as in other RNAi pathways involved in the endogenous regulation of target mRNAs [8].

Although epimutants can arise in wild-type strains, the phenomenon is enhanced by mutations in key genes of an RdRP-dependent Dicer-independent degradation mechanism for endogenous mRNA [3,9]. This could mean that either this novel RNAi pathway directly represses the epimutation machinery or that it competes for the same target mRNAs. This degradation mechanism is considered a <u>n</u>on-canonical <u>R</u>NA interference <u>p</u>athway (called NCRIP) because it does not share the canonical core RNAi machinery. Indeed, mutational analyses showed that only RdRP enzymes, but neither Dicer nor Argonaute, participate in the NCRIP pathway [10]. The cleaving activity required to degrade target mRNAs relies on a new RNase III-like protein named R3B2, which plays the primary RNase role in the NCRIP pathway. The unique role of RdRPs (RdRP1, RdRP2, and RdRP3) in RNA degradation suggests that the NCRIP mechanism represents a first evolutionary link connecting mRNA degradation and post-transcriptional gene silencing [9].

The role of NCRIP in regulating the RNAi-dependent epimutational mechanism emphasizes the intricate network of interactions among RNAi pathways in fungi. However, the actual functional role of NCRIP in cellular processes and the importance of its regulatory effects on fungal physiology are still unknown. The large number of predicted genes that might be regulated by NCRIP suggested a pleiotropic role in fungal physiology, controlling several and diverse processes. Indeed, phenotypic analysis of mutants lacking the NCRIP pathway revealed two prominent phenotypes associated with the lack of NCRIP: *in vitro* oxidative stress resistance and reduced production of zygospores during sexual development [10].

RNAi-related mechanisms are important for the maintenance of genome stability and transposon movement in other fungal pathogens such as *Cryptococcus neoformans* [11]. In this basidiomycete, the canonical RNAi machinery plays a protective role by silencing transposable elements during mating, ensuring the genomic integrity of the progeny. A recent study in *M. circinelloides* also found an essential role for the canonical RNAi core in silencing repetitive pericentric transposable elements [12]. Interestingly, analysis of genome-wide sRNA content in epimutants that were deficient in NCRIP revealed an alteration of sRNA levels derived from transposable elements [4]. These studies reinforce the hypothesis of an inhibitory function of NCRIP over the canonical pathway during the production of epimutants. Thus, NCRIP could have a role in maintaining genome integrity through its competitive regulation of the canonical RNAi in the control of transposable elements. Moreover, the resistance to oxidative stress observed *in vitro* in NCRIP deficient mutants [10] could play a specific role for survival in stressful environments, such as those related to the host-pathogen interaction. NCRIP may also be involved in pathogenesis given the high frequency of drug-resistant epimutants in mutants of this pathway, suggesting that this regulatory mechanism could be linked to virulence in *M. circinelloides*.

Here, we show a detailed functional analysis of the NCRIP pathway, addressing the functional roles that it might play in fungal biology and pathogenesis. Consequently, we studied the complex network of genes regulated by NCRIP during saprophytic growth and macrophage phagocytosis. This study identifies the complete profile of genes and functional categories regulated by NCRIP in both conditions. Interestingly, most of the fungal genes regulated by phagocytosis are under control of NCRIP, indicating that this RNAi-based mechanism is a master regulator of the response of the pathogen to phagocytosis.

## Results

### NCRIP preferentially regulates functional processes during non-stressful conditions

The higher resistance to oxidative stress of *M. circinelloides* NCRIP-deficient mutants prompted us to identify the genes controlled by this RNAi pathway in response to the oxidative burst of macrophages during phagocytosis. To this end, we performed a transcriptomic analysis of the gene expression profiles obtained from high-throughput sequencing of mRNA (RNA-seq) from spores of the wild-type strain and mutants lacking NCRIP activity (*r3b2*Δ or *rdrp1*Δ). The spores were single-cultured in rich medium L15 (saprophytic conditions), and co-cultured with the J774A.1 cell-line of mouse macrophages (1.5:1 spore‒macrophage ratio) for 5 hours to ensure that most of the spores were phagocytosed. These macrophage samples represent the closest in vitro environment to a clinical in vivo context in which the germinating spores must rapidly overcome oxidative stress to escape from the innate immune response.

Messenger RNA was isolated and deep sequenced to analyze the transcriptional response of the control wild-type samples with or without macrophages (Fig 1A, WTM or WTC, respectively), and the two mutant samples, with macrophages (Fig 1A, *r3b2*∆M and *rdrp1*∆M) or without (Fig 1A, *r3b2*∆C and *rdrp1*∆C). We performed a principal component analysis of the expression values for all genes (mean CPM > 1.0 per gene in all conditions) to further study the variability among the samples (Fig 1A). This analysis revealed that the NCRIP mutant strains, *r3b2*Δ and *rdrp1*Δ, clustered closely together and had a distinct transcriptomic profile compared to the wild-type strain growing in saprophytic conditions without macrophages. However, when the macrophages phagocytosed the spores, the mRNA repertoire of all strains formed a closer cluster and showed a more similar profile. To identify these changes in gene expression, the genetic profiles of the two mutants were compared to the wild-type strain in the presence or absence of macrophages (S1 Dataset). A threshold of a corrected p-value under 0.05 (<u>F</u>alse <u>D</u>iscovery <u>R</u>ate [FDR] of 0.05) and a log_2_ FC ≥|1.0| was selected to consider differentially expressed genes (DEGs). The deletion of either the *r3b2* or the *rdrp1* gene caused a profound variation in the mRNA profiles of *M. circinelloides*, especially when the fungus grows without macrophages (Table 1). Under these saprophytic conditions, most DEGs trend towards upregulation in the wild-type strain, as expected from the direct repressive activity of NCRIP. However, downregulation modestly prevailed in the wild-type spores phagocytosed by macrophages, suggesting repression of a few primary direct targets of NCRIP that control a vast network of secondary targets (Table 1).

**Table 1.** Differentially expressed genes in NCRIP mutant strains compared with the wild-type strain

| Culture conditions | Strain | Upregulated genes<sup>1</sup> |  | Downregulated genes<sup>2</sup> |  |
| --- | --- | --- | --- | --- | --- |
| | | # | Average $\log_2$ FC<sup>3</sup> | # | Average $\log_2$ FC<sup>3</sup> |
| L15 5h 37 °C | *r3b2*Δ | 1685 | 2.42 ± 1.36 | 850 | -1.55 ± 0.79 |
|  | *rdrp1*Δ | 1905 | 2.46 ± 1.43 | 1452 | -1.67 ± 0.79 |
| L15 5h 37 °C + Mφ | *r3b2*Δ | 80 | 1.77 ± 1.49 | 92 | -1.64 ± 0.76 |
|  | *rdrp1*Δ | 77 | 1.86 ± 1.51 | 106 | -1.61 ± 0.84 |

<sup>1</sup>FDR ≤ 0.05, $\log_2$ FC ≥ 1.0, average $\log_2$ CPM ≥ 1.0
<sup>2</sup>FDR ≤ 0.05, $\log_2$ FC ≥ -1.0, average $\log_2$ CPM ≥ 1.0
<sup>3</sup>Average of all $\log_2$ fold change values and standard deviation

**Figure 1.**
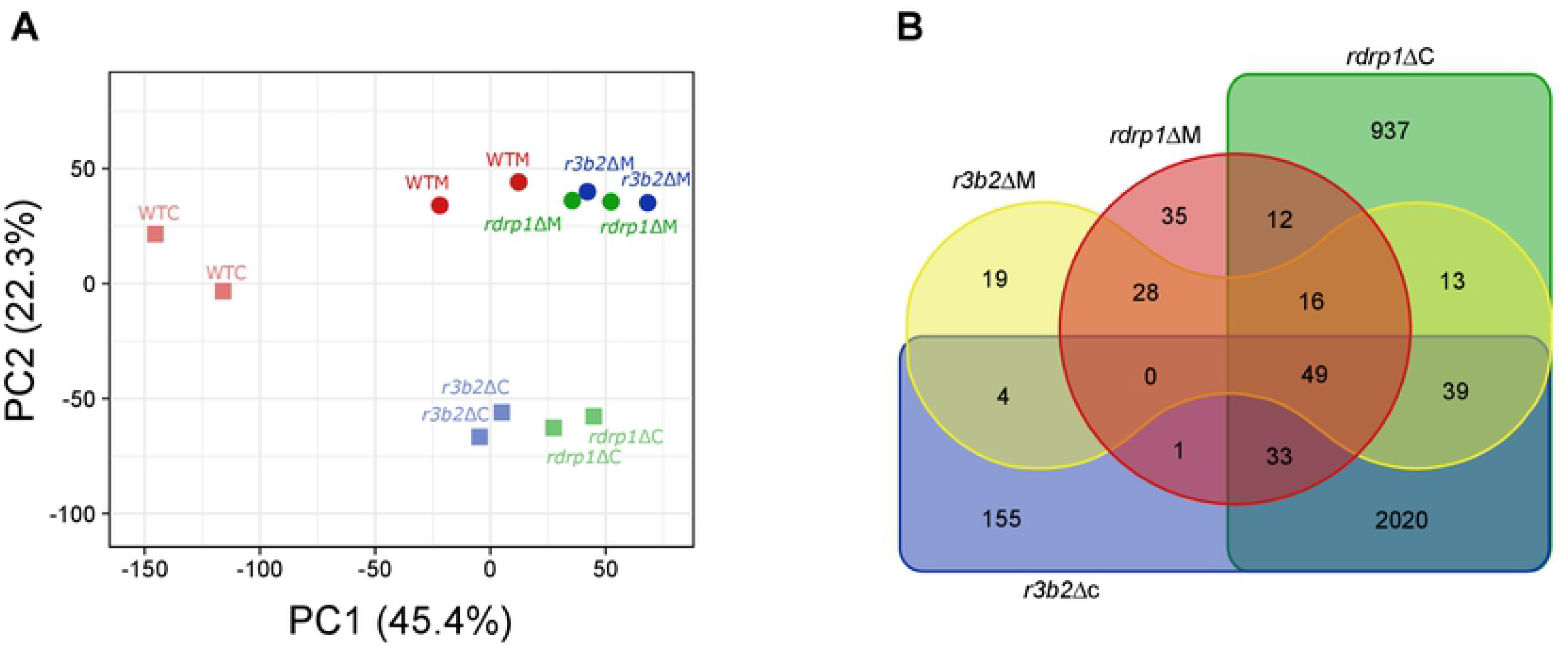
NCRIP regulates a vast gene network via the cooperation of R3B2 and RdRP1. **(A)** Principal component (PC) analysis biplot of gene abundances (in counts per million [CPM], mean CPM > 1.0), showing the similarity across the color- and shape-coded samples. **(B)** Venn diagram of significant differentially expressed genes (DEGs, log2FC ≥ |1.0|, FDR ≤ 0.05) in the depicted mutant strains and conditions compared with the wild-type strain in the same condition. DEGs overlap if they share a similar expression pattern (either up-or downregulation).

Subsequently, we searched for shared DEGs in both NCRIP mutants compared to their wild-type control, representing all comparisons in a four-way Venn diagram (Fig 1B). The analysis revealed a total of 2199 genes (> 18% of the genome) regulated by both R3B2 and RdRP1 under all conditions (Fig 1B, 28 + 16 + 13 + 0 + 49 + 39 + 1 + 33 + 2020), representing 69% of all of the DEGs. These results suggest that NCRIP regulates these genes, whereas the genes misregulated in only one or the other mutant could be the result of other independent functions of R3B2 and RdRP1. In saprophytic conditions, the two mutants showed 2141 DEGs (Fig 1B, 2020 + 39 + 49 + 33), whereas the differences were more subtle in the phagocytosed spores, and only 93 shared DEGs were identified (Fig 1B, 28 + 16 + 49). A total of 49 genes were differentially expressed in both *r3b2*Δ and *rdrp1*Δ mutants regardless of the presence or absence of macrophages, and these genes might be regulated by NCRIP and involved in essential processes required under all conditions (Fig 1B). These higher differences without macrophages and the low number of DEGs in the presence of macrophages are in accord with the results observed in the principal component analysis (Fig 1A).

To survey the possible cellular processes controlled by the NCRIP machinery in saprophytic growth without any challenge, an enrichment analysis of Eu<u>k</u>aryotic <u>O</u>rthologous <u>G</u>roups (KOG) terms was conducted. Under these non-stressful conditions, we found an enrichment in processes related to the production of extracellular structures and secondary metabolites, the remodeling of energy, amino acids, lipids and carbohydrates metabolic pathways, and by contrast an overall reduction in cytoskeletal processes (Fig 2). The genes grouped in these KOG classes indicated specific functional roles not required during phagocytosis and controlled by NCRIP. There was not any shared enrichment in any KOG class after phagocytosis of the two mutants’ spores, possibly because the gene set was too small to produce a significant result. Instead, *r3b2*Δ and *rdrp1*Δ mutants showed independent roles in amino acid transport and metabolism, and chromatin structure and dynamics, respectively, suggesting that these genes perform specific roles required during phagocytosis that are not controlled by NCRIP. Altogether, these results indicated a preferential activity of NCRIP under non-stressful conditions, when the spores are cultured without macrophages.

**Figure 2.**
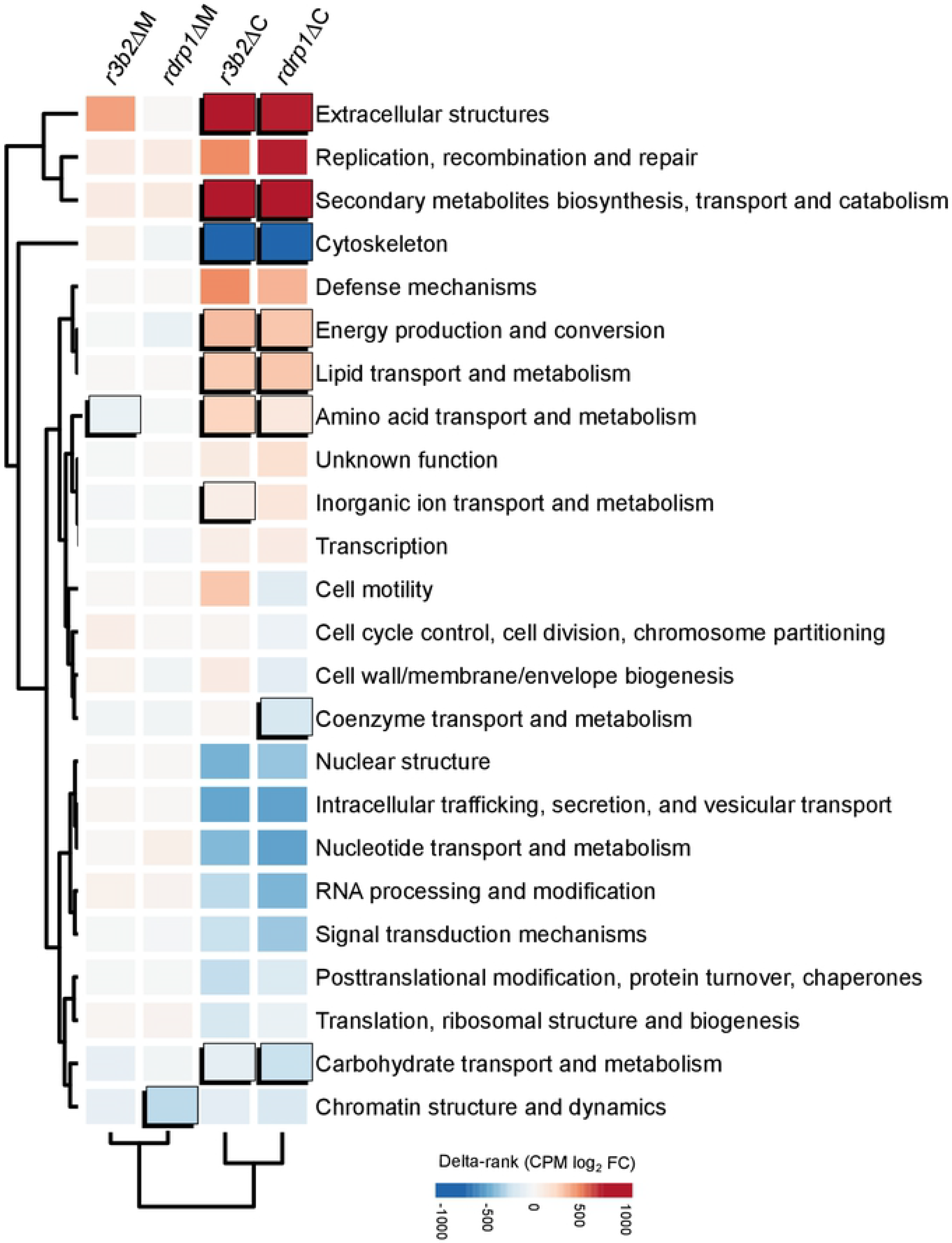
NCRIP regulates key functional categories involved in saprophytic growth. Enrichment analysis of DEGs in each Eu<u>k</u>aryotic <u>O</u>rthologous <u>G</u>roups (KOG) class. Significant enrichments (Fisher’s exact test, P ≤ 0.05) in a given mutant strain and condition compared to the wild-type strain is shown as an uplifted rectangle. A measure of up-(red) or downregulation (blue) of each KOG class is represented as a colored scale of delta-rank values (the difference between the mean differential expression value of all genes in each KOG class and the mean differential expression value of all other genes). KOG classes and experimental conditions (mutant strains and presence/absence of macrophages) are clustered according to the similarity of their delta rank values.

### NCRIP repressed the genetic response to phagocytosis during non-stress conditions

Previous studies revealed an intricate network of genes activated in response to phagocytosis, which is essential for the pathogenic potential of Mucorales [13]. Considering the large number of genes regulated by NCRIP, and the functional processes involved, we postulated that some of these genes might participate in the response to phagocytosis. To address this hypothesis, we analyzed the DEGs detected in response to phagocytosis in the wild-type strain and the *rdrp1*Δ and *r3b2*Δ mutants and presented the results in a three-way Venn’s diagram (Fig 3A). The most marked result from this analysis is the high number of genes (a total of 908 out of 1156) differentially expressed only in the wild type during phagocytosis, but not in the *rdrp1*Δ or *r3b2*Δ mutant. Therefore, these results identified a broad set of genes responding to phagocytosis in the wild-type strain that is unable to respond in the mutants lacking the NCRIP pathway. Two alternative possibilities could explain these results: either these genes required a functional NCRIP for their activation during phagocytosis or NCRIP is repressing them under non-stressful conditions without macrophages.

**Figure 3.**
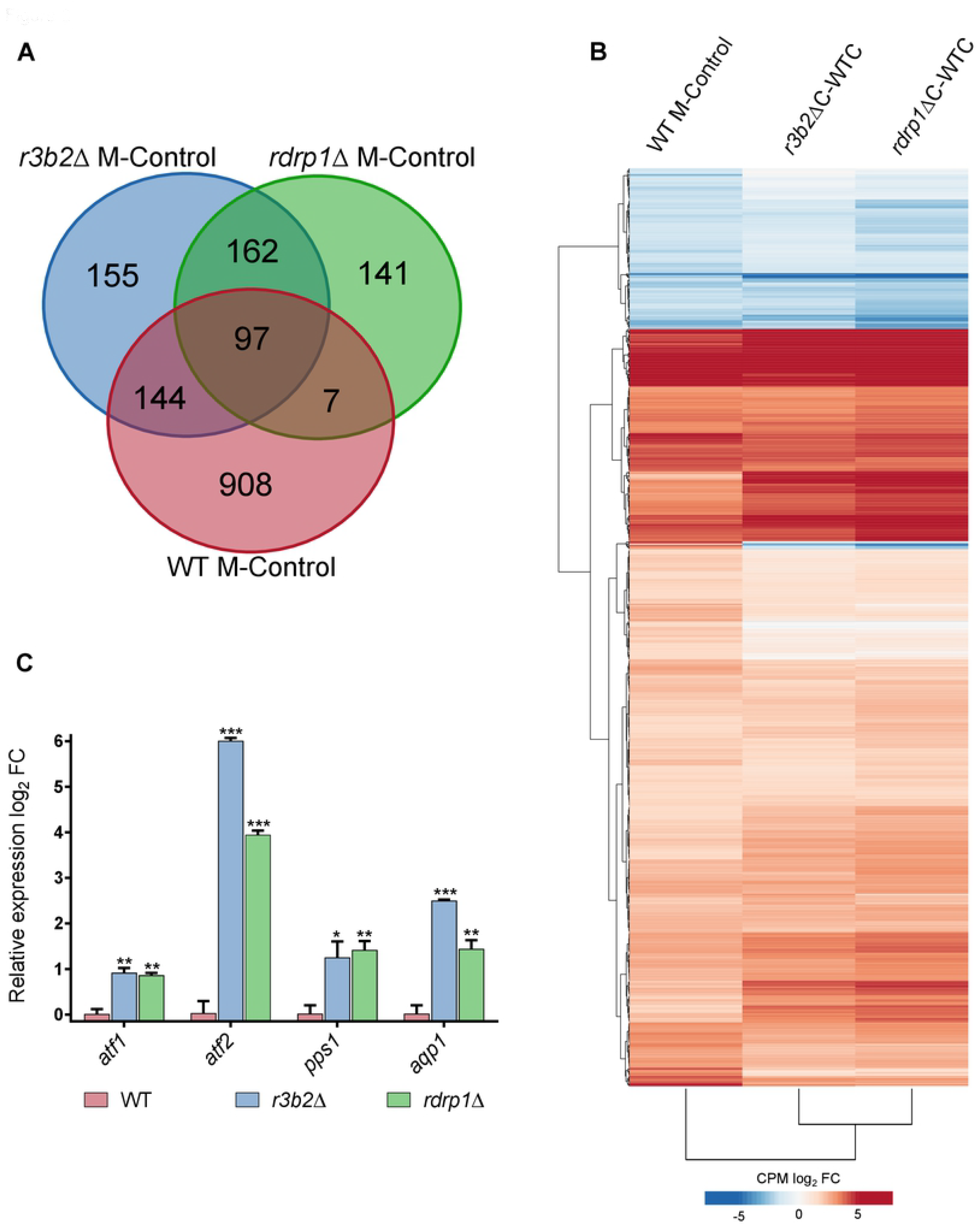
NCRIP controls the response to macrophage phagocytosis by inhibiting its targets under non-stressful conditions. **(A)** Venn diagram of DEGs (log2FC ≥ |1.5|, FDR ≤ 0.05) in the wild-type, *r3b2*Δ, and *rdrp1*Δ mutant strains phagocytosed by macrophages compared with their non-phagocytosed controls. DEGs overlap if they share a similar expression pattern (either up- or downregulation). **(B)** Heatmap of the 908 DEGs found in (A) that were responding to phagocytosis in the wild-type strain but were not differentially expressed in either NCRIP mutant strain. Genes and experimental conditions are clustered by similarity to compare the response to phagocytosis in the wild-type strain with the response of the NCRIP mutants in non-stressful conditions. DEGs are color-coded to depict the degree of upregulation (red) or downregulation (blue) in each condition. **(C)** Bar plot of *atf1*, *atf2*, *pps1* and *aqp1* expression differences in R3B2Δ, and RdRP1Δ mutant strains compared with the wild-type strain in non-stressful conditions, i.e. incubation in cell-culture medium without macrophages for 5 hours. Log_2_ fold-change differential expression levels were quantified by RT-qPCR and normalized using *rRNA* 18S as an internal control. Error bars correspond to the SD of technical triplicates and significant differences are denoted by asterisks (* for P ≤ 0.05, ** for P ≤ 0.005, and *** for P < 0.0001 in an unpaired t-test).

To clarify the role of NCRIP in the regulation of this gene network, we further analyzed their expression levels in three different comparisons: WTM vs. WTC, *r3b2*∆C vs. WTC, and *rdrp1*∆C vs. WTC (Fig 3B). Surprisingly, the differential expression in both *rdrp1*Δ and *r3b2*Δ mutants compared to the wild-type strain in saprophytic conditions was almost identical to those found in the wild-type strain responding to phagocytosis (Fig 3B). This analysis showed a group of genes activated both by macrophage-mediated phagocytosis in the wild-type strain and by the lack of *rdrp1* or *r3b2* (Fig 3B, coincidences in red). This group of activated genes may correspond to primary target genes repressed by NCRIP, suggesting a negative regulation of NCRIP in the absence of macrophages that is released upon phagocytosis in the wild-type strain. A second group consists of genes repressed both by the presence of macrophages in the wild-type strain and the lack of *rdrp1* or *r3b2* (Fig 3B, coincidences in blue). These genes could be acting as secondary targets of the primary gene set.

Previous studies identified gene expression profiles during the phagocytosis of *M. circinelloides*’ wild-type spores [13]. Those profiles were validated by quantitative RT-PCR using the following representative marker genes: *atf1, atf2, pps1*, and *aqp1*. These marker genes showed a significant induction during macrophage phagocytosis and are essential for this fungal pathogen to survive and cause infection. Our transcriptomic analysis found that all of these marker genes were also controlled by NCRIP during saprophytic growth (S1 Dataset), and thus, they were employed here to validate the transcriptional pre-activation observed in the mutants *rdrp1*Δ and *r3b2*Δ without macrophages (Fig 3C). We found that the four marker genes showed significant induction in the two mutants without macrophages, similar to the previously reported increased expression observed in the wildtype during phagocytosis [13], indicating that NCRIP controls the response to phagocytosis by repressing it during non-stressful conditions.

Functional enrichment analysis of this gene set was performed to further understand the biological processes controlled by this response (Fig 4). We observed a clear alteration of the metabolism and transport that affects both carbohydrates and lipids because the corresponding KOG classes included genes down- and upregulated. This was associated with an overrepresentation of upregulated genes involved in amino acid transport and metabolism, suggesting a metabolic change linked to the germination process inside the phagosome. The harsh phagosomal environment might also be responsible for the induction of genes related to the biosynthesis, transport, and catabolism of secondary metabolites and extracellular structures to defend the fungus from an oxidative challenge. Enrichment of downregulated genes involved in cell motility and cytoskeletal processes could also be a part of a fungal strategy to produce competent extracellular structures needed for survival and germination inside the phagosome. This response was functionally similar to that observed from the whole gene profile identified in the NCRIP mutants under non-stressful conditions (Fig 2).

**Figure 4.**
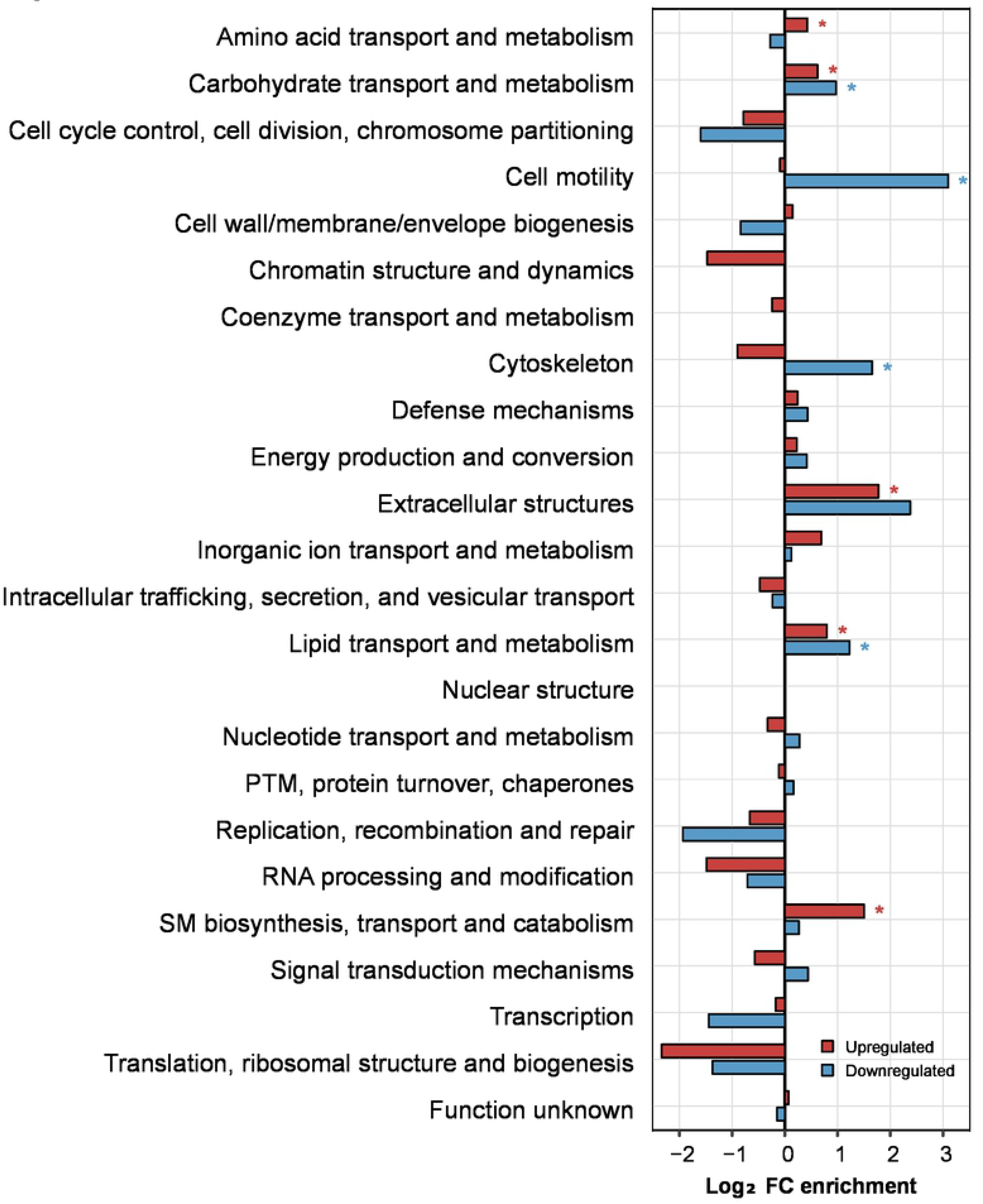
Functional analysis of the response controlled by NCRIP during macrophage interaction. Enrichment analysis of the 908 DEGs controlled by NCRIP, showing up-(red) and downregulated (blue) genes. Fold-enrichment in each KOG category is plotted and significant enrichments (Fisher’s exact test, P ≤ 0.05) is marked by an asterisk (*).

### NCRIP negatively regulates the protective role of the canonical RNAi pathway in the suppression of Grem-LINE1s retrotransposons

The results presented above indicate that NCRIP represses genes during non-stressful conditions and then releases its control upon phagocytosis by macrophages, a clear challenging stimulus. However, the expression profiles analyzed did not reveal any specific pathway involved in sporulation or mating. Because mutants in the machinery of NCRIP also display defects in the production of zygospores during mating, we hypothesized that NCRIP might also contribute in the regulation of genes involved in other stresses such as antifungal agents [4,9] and genomic integrity stress, which could alter complex cell processes involved in mating [10]. A recent study supported this hypothesis, unveiling a direct link between the canonical RNAi pathway and the protection of genome integrity against transposable elements in *M. circinelloides* [12]. The pericentric regions of *M. circinelloides* contain a large number of L1-like retrotransposable elements of the Mucoromycotina species called Grem-LINE1s, which are actively silenced by the canonical RNAi machinery. As suggested previously [9], NCRIP might regulate the RNAi canonical core during the control of these transposable elements by suppressing the epimutational pathway. In this sense, we characterized the production of siRNAs from Grem-LINE1 transcripts in the pericentric regions of the wild-type strains and the *r3b2*Δ *and rdrp1*Δ mutants (Fig 5A). The pericentric regions are almost depleted of siRNAs in the *ago1*Δ and *dcl1 dcl2*Δ mutants, whereas the wild-type strain exhibited an active production of siRNAs aligned to these loci, as previously reported [12]. Interestingly, the *r3b2*Δ *and rdrp1*Δ mutants displayed an exaggerated production of siRNAs compared to the wild-type strain, especially targeting the second open reading frame (ORF2) and its reverse transcriptase domain (RVT). The over-accumulation of siRNAs (≥ 1.5 log_2_ FC) is consistent among all Grem-LINE1s in the *r3b2*Δ *and rdrp1*Δ mutants (Fig 5B). These results suggest an enhanced activity of the canonical RNAi machinery degrading the target retrotransposons when NCRIP is not active and therefore, a negative regulatory role for NCRIP.

**Figure 5.**
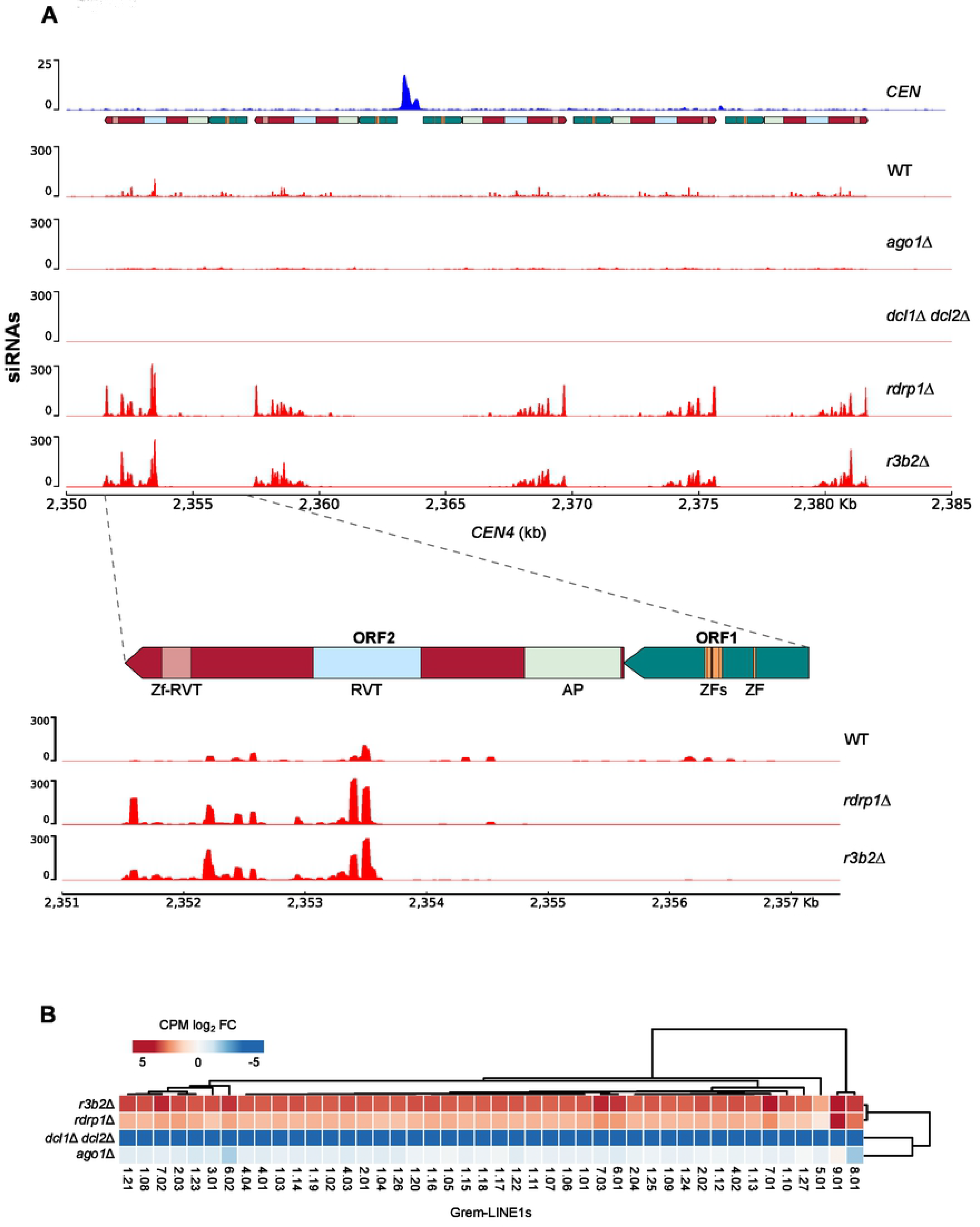
NCRIP competes with the epimutational pathway to regulate transposable elements. **(A)** Genomic view of centromeric chromatin (*CEN4*) displaying the kinetochore-binding region enrichment that marks the centromere (*CEN*, blue), annotation of transposable elements (colored blocks), and transcriptomic data of sRNAs (red) in *M. circinelloides* wild-type, epimutational pathway (*ago1∆*, double *dcl1∆/dcl2∆*) and NCRIP (*rdrp1∆* and *r3b2∆*) deletion mutant strains after 48 h of growth in rich medium. The zoom below represents a single Grem-LINE1 and the NCRIP regulation by small RNAs. Open reading frames (ORF1 in green arrows and ORF2 in red arrows) and protein domains predicted from their coding sequences are shown as colored blocks (zf-RVT, zinc-binding in reverse transcriptase [PF13966]; RVT, reverse transcriptase [PF00078]; AP, AP endonuclease [PTHR22748]; and ZF, zinc finger [PF00098 and PF16588]). **(B)** Heatmap of the differential expression of <u>G</u>enomic retrotransposable <u>e</u>lements of <u>M</u>ucoromycotina LINE1-like (Grem-LINE1s) in the depicted RNAi mutants compared to the RNAi-proficient wild-type strain. Grem-LINE1s are numbered according to Navarro et al. classification [12].

### Lack of NCRIP decreases virulence

Virulence is a complex trait that depends on multiple genes and is controlled by different biological processes [14]. The fact that NCRIP regulates the expression of hundreds of genes involved in the response to phagocytosis suggests a role in controlling critical pathways involved in virulence. To test this hypothesis, the *r3b2*Δ and *rdrp1*Δ mutants were used to perform survival assays in an immunosuppressed mouse model, previously validated as a host model for infections with *M. circinelloides* [15]. The survival rates were compared to those of mice injected with the wild-type virulent strain R7B and the NRRL3631 strain, a natural soil isolate that served as an avirulent mock control [16]. The results of these assays showed a significant reduction in virulence of the two mutant strains (Log-rank Mantel-Cox test, p = 0.0061 in *rdrp1*Δ vs. R7B; p = 0.040 in *r3b2*Δ vs. R7B; Fig 6), indicating that NCRIP controls the expression of genes involved in virulence.

**Figure 6.**
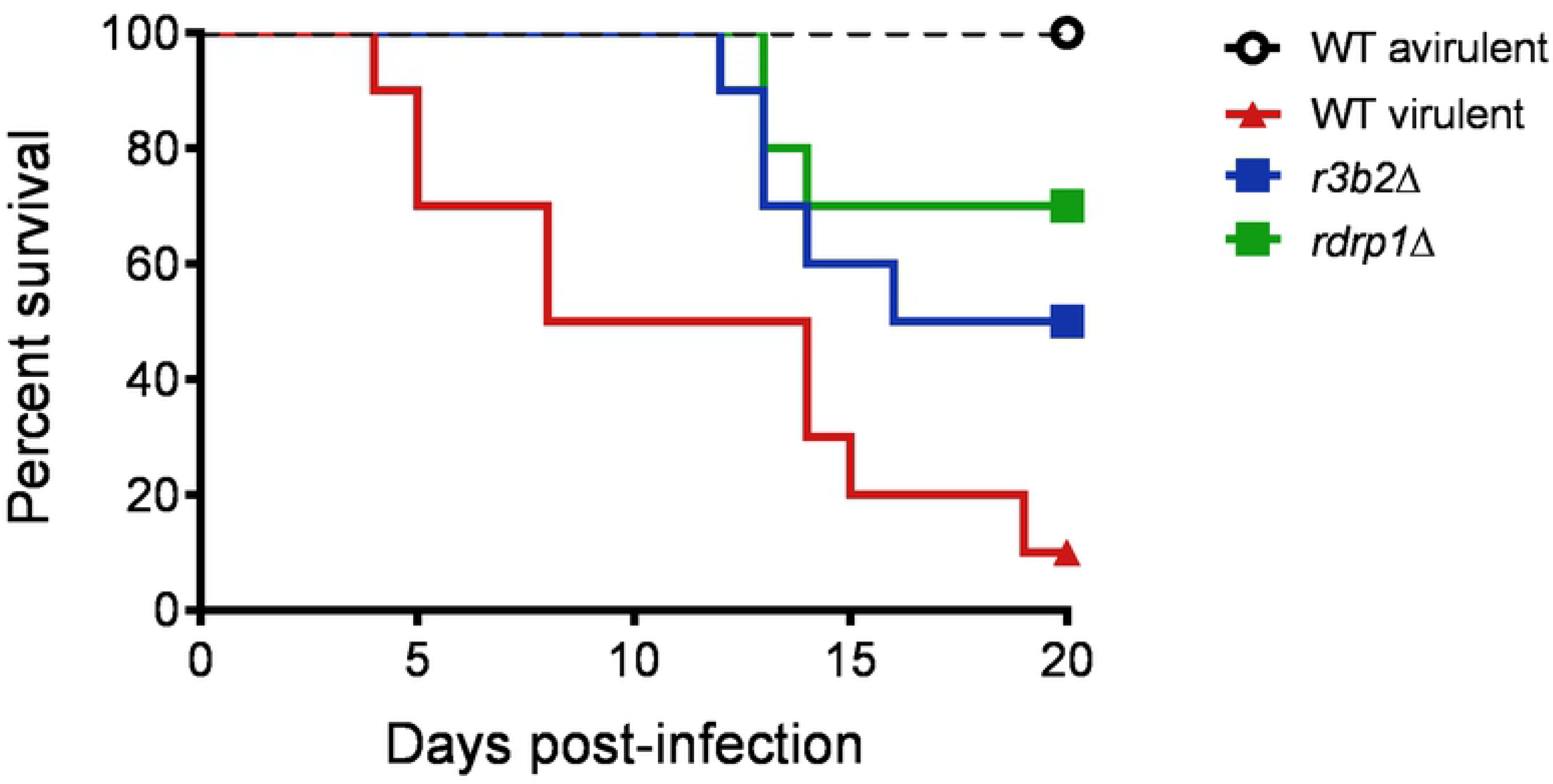
NCRIP is involved in mucormycosis. The virulence of *r3b2*Δ and *rdrp1*Δ mutant strains was assessed in a survival assay using immunosuppressed mice as a mucormycosis model. Groups of ten mice were infected intravenously with 1×10^6^ spores from each strain (color-coded). Survival rates were statistically analyzed for significant differences (* for P ≤ 0.05 in a Mantel-Cox test) compared with a virulent control strain. NRRL3631 was used as an avirulent mock control of infection.

## Discussion

Among the diversity of RNAi pathways in *M. circinelloides*, NCRIP is the most recently discovered. It is a new mechanism that remains largely uncharacterized, and its functional role in fungal physiology is the central unanswered question. Is it a non-canonical RNAi degradation mechanism that clears and turns over damaged RNAs? Or does it play a regulatory function controlling the expression of mRNAs at specific levels depending on cellular requirements? The results obtained in this study unveiled a complex regulatory role of NCRIP in fungal physiology rather than a simple degradation mechanism for functional or damaged RNAs. Thus, we identified hundreds of genes regulated by NCRIP, including genes involved in survival during phagocytosis. The analysis of the spore response to the phagosome revealed a derepression of a complex gene network activated in the fungal spore after the interaction with macrophages. Moreover, we identified a negative regulatory role of NCRIP over the canonical RNAi pathway in the control of transposable elements, extending the functional complexity of this mechanism beyond the control of cellular mRNA levels. These complex functional roles of NCRIP correlated with the pleiotropic phenotypes observed in mutants of this pathway, including the reduced virulence described here.

Regarding the gene network regulated by NCRIP, previous studies suggested a broad regulatory function of this pathway based on the discovery of 611 loci producing sRNAs in a *dicer*-independent *rdrp*-dependent manner [10]. Here, we have directly analyzed the transcriptomic profiles in NCRIP key mutants, identifying a substantial number of DEGs in both *rdrp1* and *r3b2* mutants compared to the wild-type strain. However, a significantly lower number of genes were regulated in these mutants upon phagocytosis compared to the complex response observed in the wild-type strain [13]. The principal component analysis and the comparison of the four profiles among them further confirmed this strong bias among samples. Thus, NCRIP showed a differential regulatory intensity when rich and stressful environments were compared. These results prompted us to hypothesize a repressive regulatory role of NCRIP under non-stress conditions; hence, upon cellular challenges (like drug exposure or phagocytosis), the repression would cease allowing the activation of the corresponding gene response. Previous studies support this hypothesis, finding a similar regulatory mechanism in the epimutational pathway, which also suggested a negative regulatory role under no stress conditions [9]. A more in-depth analysis of the gene profiles and their expression levels supports our hypothesis because the mutants activated the gene response to phagocytosis before the interaction with macrophages. These findings might explain the augmented oxidative stress resistance observed in the NCRIP mutants in vitro [10]. Thus, the mutants in NCRIP could be unable to respond properly to stress because they show constitutive expression of these genes which might lead to a pre-exposure adaptation to the stimuli.

Another regulatory role of NCRIP identified in this work unveiled a novel genetic mechanism in which the canonical RNAi pathway and NCRIP work as antagonistic dual machinery to control the movement of transposable elements. Previous studies reported that most sRNAs produced by the NCRIP machinery map to exonic regions, whereas a minimal amount of these sRNAs were found in intergenic regions and transposable elements [10]. However, these studies were developed using initial annotation versions of the *M. circinelloides* genome before the identification of the centromeric regions. Once the centromeric regions were assembled, they were further characterized as rich in repetitive sequences and Grem-LINE1 retrotransposons [12]. The expression of the mobile elements is suppressed by the canonical RNAi machinery and correlates with an abundant production of sRNAs and low mRNA levels. In this work, we found an exacerbated production of sRNAs from centromeric transposons in the NCRIP mutants, indicating an overactivation/derepression of the canonical RNAi pathway. These results suggest a negative regulatory role of NCRIP over the canonical RNAi pathway, analogous to the inhibitory role the NCRIP pathway exerts over the epimutational pathway. Via the canonical RNAi machinery, the epimutational pathway silences target genes to overcome growth inhibition caused by antifungal compounds and generates epimutant strains that are resistant to drugs [3,4]. Conversely, the inactivation of NCRIP leads to an overproduction of epimutant strains [9], suggesting either a competition between NCRIP and the epimutational pathway for the transcripts of the target gene or repression of NCRIP over the canonical mechanism. Here, we propose that the same mechanism is operating in the control of the movement of pericentromeric retrotransposons. This hypothesis explains why when the NCRIP is inactive most of the retrotransposons are controlled by the canonical RNAi machinery and there is enhanced production of sRNAs in *r3b2*Δ and *rdrp1*Δ mutants. Thus, both interacting RNAi pathways could be essential for genome stability and integrity.

The role of the RNAi machinery in protecting genome integrity against the movement of transposons is important during mating in several fungal models. In *C. neoformans*, the mechanism of sex-induced silencing (SIS) defends the genome against transposons during sexual development, whereas in several ascomycetes [17–19] an RNAi mechanism operates to silence unpaired DNA in meiosis, including transposons [20]. These surveillance mechanisms that protect genome integrity rely on the RNAi canonical core, as in *M. circinelloides*. However, in this fungus, the canonical RNAi pathway coexists with a regulatory mechanism based on NCRIP, which has not been described in other fungal groups [10]. It is tempting to speculate that both the canonical mechanism and NCRIP perform a fine control over retrotransposable movements to gain genetic diversity in particular stressful conditions, allowing a transient activation of the retrotransposons to overcome the insult. Alteration of this precise control may be responsible for the defective mating observed in NCRIP mutants.

The pre-activated state of NCRIP mutants and their previously described oxidative stress resistance suggests an advantage to resist the oxidative attack of macrophages. However, our results showed a reduced pathogenic potential in both *rdrp1*Δ and *r3b2*Δ mutants, indicating that NCRIP is necessary for virulence in Mucorales. These results reveal that the resistance to oxidative stress in vitro did not improve the pathogenicity of the mutants during in vivo interactions. The complex environment of the host during the initial steps of phagocytosis could explain these results, because the fungus must respond to phagosomal conditions, including oxidative stress, nutritional starvation, and pH acidification. The intricate transcriptomic response displayed by the spores to counteract the host was not fully replicated in the NCRIP mutants, and these mutants show DEGs that are not regulated by phagocytosis (Fig 3A). On the other hand, the genetic deregulation in NCRIP mutants might affect other fungal responses required during the response to phagocytosis, or in further infection steps, such as tissue invasion, resulting in a final negative balance for the fungal spore.

Our functional study unveiled a complex gene network conditionally regulated by NCRIP. The analysis of this gene network revealed a remarkable function of NCRIP in the negative regulation of the genetic response elicited during phagocytosis, suggesting an essential role for this pathway in host-pathogen interactions. Altogether, the identification of a large number of genes regulated by NCRIP and the subset involved in the response to phagocytosis confirm the broad regulatory role of NCRIP, arguing against a simpler role in clearance and turnover of RNAs. Instead, NCRIP emerges as a mechanism controlling an extensive network of genes involved in different cellular processes, with the capability of regulating them differentially after environmental challenges that include antifungals agents, phagocytosis, and virulence. The role of NCRIP controlling the genetic response to phagocytosis and the final phenotypic balance impairing virulence are new contributions to understanding the difficult to treat and challenging to manage infection of mucormycosis.

## Materials and Methods

### Fungal strains, cell cultures, and RNA purification

The fungal strains used in this work derived from *M. circinelloides* f. *lusitanicus* CBS277.49. The wild-type control strain for the RNA-seq analysis and virulence assays is R7B [21]. The strains defective in the NCRIP are MU419 (*rdrp1*∆) [22] and MU412 (*r3b2*∆) [10]. All the strains compared for the gene expression analysis were auxotroph for leucine. The strain NRRL3631 was used as an avirulent control for the mice infection experiments [23]. *M. circinelloides* cultures were grown in rich media YPG pH 4.5 at 26ºC for optimal growth and sporulation. Spores were harvested and filtered using a Falcon® 70 µm cell strainer before confronting with macrophages or animal models. The host-pathogen interactions were performed confronting spores from R7B, MU419, and MU412 with mouse macrophages (cell line J774A.1; ATCC TIB-67) in a ratio 1.5:1 (spores:macrophages) following the protocol described in [13]. In summary, the interactions were maintained at 37ºC in L15 medium (Capricorn Scientific) supplemented with 20% of Fetal Bovine Serum (Capricorn Scientific) for 5 hours, ensuring all the spores were phagocytosed. For saprophytic conditions, the same concentration of spores was cultured in L15 medium as described.

For RNA purification, two replicates of each sample were pooled, and RNA was extracted using the RNeasy plant minikit (Qiagen, Hilden, Germany), following the manufacturer procedure.

### RNA-sequencing analysis for gene expression and small RNA production

Raw datasets were quality-checked using FASTQC v0.11.8 before and after removing adapter and contaminant sequences with Trim Galore! v.0.6.2 (http://www.bioinformatics.babraham.ac.uk/projects/). Messenger RNA reads were aligned to the *M. circinelloides* f. *lusitanicus* v2.0 genome (Mucci2 [24]) using STAR v.2.7.1a [25] and the subsequent <u>B</u>inary <u>A</u>lignment <u>M</u>aps (BAM) were used to create individual count matrices with HTSeq v.0.9.1 [26], excluding multi-mapping reads. Differential gene expression and <u>p</u>rincipal <u>c</u>omponents <u>a</u>nalysis (PCA) were performed by *limma* package v.3.38.3 [27]; genes above a reliable threshold used in a previous study [13] (False Discovery rate [FDR] ≤0.05; log_2_fold change [log_2_ FC] ≥1.0; and average <u>c</u>ount <u>p</u>er <u>m</u>illion reads (CPM) ≥1.0) were considered differentially expressed genes (DEGs) and used for downstream analyses unless more stringent criteria is stated. DEGs were classified according to Eu<u>k</u>aryotic <u>O</u>rthologous <u>G</u>roups (KOG) and <u>G</u>ene <u>O</u>ntology (GO) terms using EggNOG-mapper v2.0 [28,29] to perform KOG class enrichment analyses with KOGMWU package v.1.2 [30]; statistical significance was assessed with a Fisher’s exact test (p-value ≤0.05). Small RNA reads were obtained from previous studies (see Data Availability) and aligned to the *M. circinelloides* f. *lusitanicus* MU402 v1.0 genome (Muccir1_3 [12]) using the <u>B</u>urrows-<u>W</u>heeler <u>A</u>ligner (BWA) v.0.7.8 [31]. This genome was PacBio-sequenced using long reads and thus, exhibit a greater content of repeated elements [12]. The number of overlapping aligned reads per 25-bp bin was used as a measure of coverage, obtaining Coverage was normalized to <u>b</u>ins <u>p</u>er <u>m</u>illion reads (BPM) in 25-bp bins with deepTools v3.2.1 [32] bamCoverage function. The resulting bigWig files were visualized with the deepTools pyGenomeTracks module using the centromeric and transposable element annotations found by Navarro-Mendoza et al. 2019 [12].

### RT-qPCR quantification

Replicate samples for the host-pathogen interactions and control conditions were used for RT-qPCR analysis. Once the mRNA was purified and treated with TURBO DNase (Thermo Fisher), the cDNA was synthesized from 1µg of total RNA using the iScript cDNA synthesis kit (Bio-Rad). The RT-qPCR was performed in triplicate using 2X SYBR green PCR master mix (Applied Biosystems) with a QuantStudio TM 5 flex system (Applied Biosystems) using 2X SYBR green PCR master mix (Applied Biosystems) following the supplier’s recommendations. To ensure non-specific amplification, non-template control and melting curve were tested. The primer sequences used for the quantification of genes *atf1*, *atf2*, *pps1*, *aqp1*, and rRNA *18S* are listed in S1 Table. The efficiencies of every pair of primers were approximately identical; thus the relative gene expression of the target genes was obtained the delta-delta cycle threshold (ΔΔCt) method, normalizing for the endogenous control rRNA *18S*.

### Virulence assays

The murine infection assays for Mucorales virulence were performed using OF-1 male mice weighing 30g (Charles River, Barcelona, Spain) [13,23,33]. The mice were immunosuppressed with the administration of cyclophosphamide (200 mg/kg of body weight) via intraperitoneal, 2 days prior to infection and once every 5 days thereafter. Groups of 10 mice were challenged with 1×10^6^ spores of the strains R7B, NRRL3631, MU419 and MU412. The infections were performed intravenously via retroorbital injection following the protocol described by Chang et al. 2019 [5]. Before the injection, mice were anesthetized by inhalation of isoflurane, and then the animals were visually monitored while recovering from the anesthesia. Mice were housed under established conditions with free food and autoclaved water. The animal welfare was checked twice daily for 20 days, and those following the criteria for discomfort were euthanized by CO_2_ inhalation. The significance of survival rates was quantified using the Kaplan-Meier estimator (Graph Pad Prism). Differences were considered statistically significant at a p-value of ≤0.05 in a Mantel-Cox test.

### Data availability

The raw data and processed files generated by this work are deposited at the <u>G</u>ene <u>E</u>xpression <u>O</u>mnibus (GEO) repository and are publicly available through the project accession number GSE142543. These data were compared to a wild-type strain in the same conditions, previously available at GEO [13] under the following sample accession numbers: GSM3293661 and GSM3293662 (wild-type strain single-cultured); and GSM3293663 and GSM3293664 (wild-type strain co-cultured with mouse macrophages). Mucci2 [24] and Muccir1_3 [12] genomes and annotation files can be accessed at the Joint Genome Institute (JGI) website (http://genome.jgi.doe.gov/) and used under the JGI Data Usage Policy. The small RNA raw data were available to the public through the following NCBI SRA run accession numbers: SRR039123 (wild-type strain) [34], SRR836082 (*ago1*∆ mutant strain) [35], SRR039128 (double *dcl1*∆ *dcl2*∆ mutant strain) [34], SRR039126 (*rdrp1*∆ mutant strain) [34], and SRR1576768 (*r3b2*∆ mutant strain) [10].

### Ethics statement

To guarantee the welfare of the animals and the ethics of any procedure related to animal experimentation all the experiments performed in this work complied with the Guidelines of the European Union Council (Directive 2010/63/EU) and the Spanish RD 53/2013. Experiments and procedures were supervised and approved by the University of Murcia Animal Welfare and Ethics Committee and the Council of Water, Agriculture, Farming, Fishing and Environment of Murcia (Consejería de Agua, Agricultura, Ganadería, Pesca y Medio Ambiente de la CARM), Spain (authorization number REGA ES300305440012).

## Acknowledgments

This investigation was supported by the Ministerio de Economía y Competitividad, Spain (BFU2015-65501-P, co-financed by FEDER, and RYC-2014-15844) and the Ministerio de Ciencia, Innovación y Universidades, Spain (PGC2018-097452-B-I00, co-financed by FEDER). C.P.-A. and M.I.N.-M. were supported by predoctoral fellowships from the Ministerio de Educación, Cultura y Deporte, Spain (FPU14/01983 and FPU14/01832, respectively). We thank Joseph Heitman for critical reading of the manuscript.

## Supporting information

**S1 Dataset. Differentially expressed genes in NCRIP mutants.**

**S1 Table. Primers used in the study.**

